# The evolutionary origins of primate scleral coloration

**DOI:** 10.1101/2021.07.25.453695

**Authors:** Alex S. Mearing, Judith M. Burkart, Jacob Dunn, Sally E. Street, Kathelijne Koops

## Abstract

Primate gaze following behaviors are of great interest to evolutionary scientists studying social cognition. The ability of an organism to determine a conspecific’s likely intentions from their gaze direction may confer an advantage to individuals in a social group. This advantage could be cooperative and/or competitive. Humans are unusual in possessing depigmented sclerae whereas most other extant primates, including the closely related chimpanzee, possess dark scleral pigment. The origins of divergent scleral morphologies are currently unclear, though human white sclerae are often assumed to underlie our hyper-cooperative behaviors. Here, we use phylogenetic generalized least squares (PGLS) analyses with previously generated species-level scores of proactive prosociality, social tolerance (both n=15 primate species), and conspecific lethal aggression (n=108 primate species) to provide the first quantitative, comparative test of three complementary hypotheses. The cooperative eye [M. Tomasello, B. Hare, H. Lehmann, J. Call, J. Hum. Evol. 52, 314–320 (2007)] and self-domestication [B. Hare, Annu. Rev. Psychol. 68, 155-186 (2017)] explanations predict white sclerae to be associated with cooperative, rather than competitive, environments. The gaze camouflage hypothesis [H. Kobayashi, S. Kohshima, J. Hum. Evol. 40, 419-435 (2001)] predicts that dark scleral pigment functions as gaze direction camouflage in competitive social environments. We show that white sclerae in primates are associated with increased cooperative behaviors whereas dark sclerae are associated with reduced cooperative behaviors and increased intra-specific lethal aggression. Our results lend support to all three hypotheses of scleral evolution, suggesting that primate scleral morphologies evolve in relation to variation in social environment.

## Introduction

The primate order contains a remarkable amount of variation in external ocular morphology (**Figure 1**), including differences in scleral volume, width-height ratios and pigment profiles (1-7). The former two measurements have been linked in phylogenetic comparative analyses to social (i.e., group size and neocortex ratio), ecological (i.e., habitat use) and life history (i.e., body mass) drivers (3). However, to our knowledge, no comparative quantitative study has yet examined the relationship between ocular pigment and social behavior across primate species. Humans are often considered to possess unique ocular configurations (1-3, 6-7). We possess especially large width to height ratios, especially large scleral volumes (1-4) and white, depigmented sclerae (1-2, 5-7). Contrastingly, most non-human primate species (hereafter ‘primates’), including the closely related chimpanzee, instead synthesize dark scleral pigment (1-2, 6-7). Interestingly, the equally closely related bonobo possesses sclerae of an intermediate average brightness between humans and chimpanzees (5).

**Figure 1.**
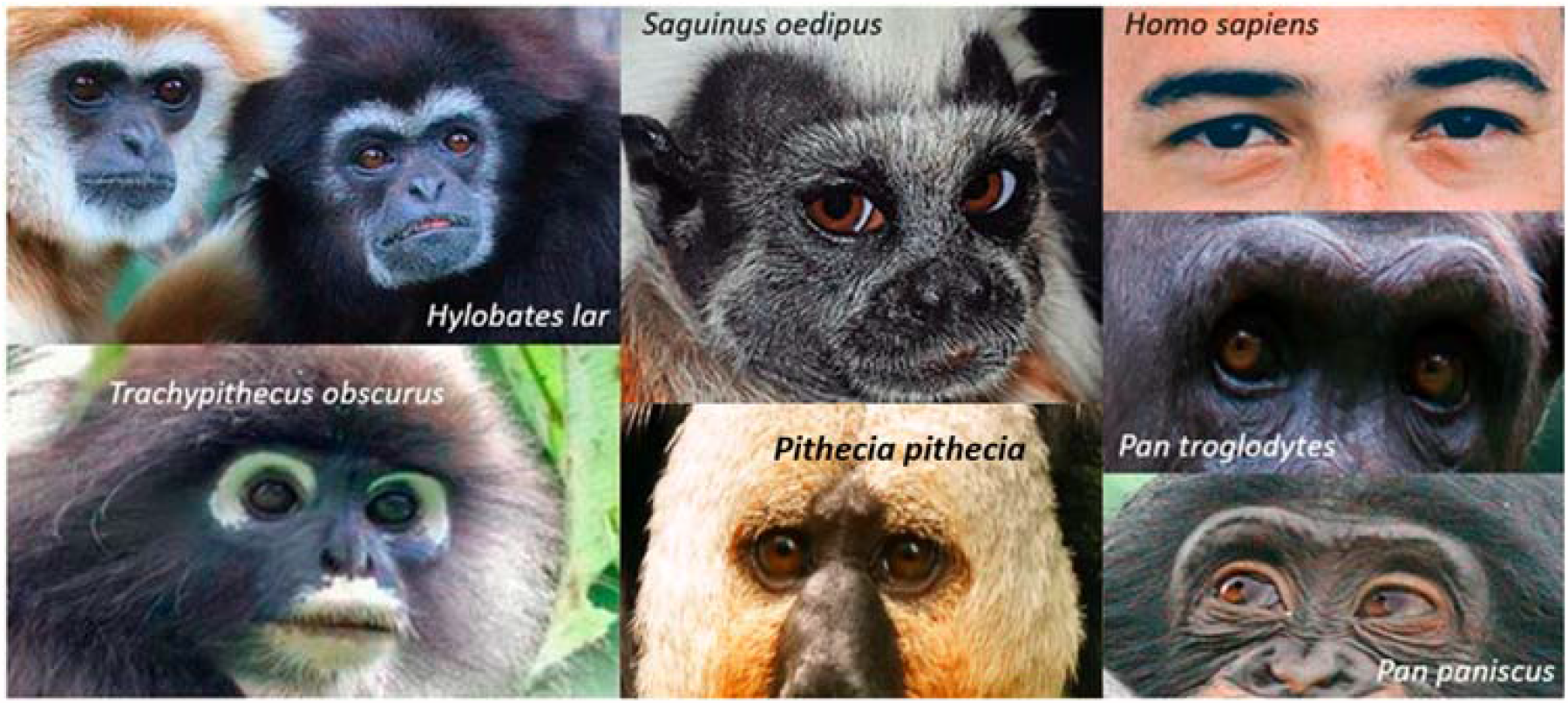
Ocular diversity in the primate order. All photos under creative commons license unless otherwise specified. H. lar credit to user: MatthiasKabel (https://commons.wikimedia.org/wiki/Hylobatidae#/media/File:Hylobates_lar_pair_of_white_and_black_02.jpg). T. obscurus credit to Lip Kee Yap (https://commons.wikimedia.org/wiki/File:Trachypithecus_obscurus.jpg). S. oedipus credit to Michael Gäbler (https://en.m.wikipedia.org/wiki/File:Saguinus_oedipus_(Linnaeus,_1758).jpg). P. pithecia credit to Hans Hillewaert (https://en.wikipedia.org/wiki/Pitheciidae#/media/File:Pithecia_pithecia.jpg). H. sapiens photo is in the public domain, credit to Fernanda Latronica (https://www.pexels.com/photo/close-up-photography-of-bearded-man-713520/). P. troglodytes photo provided by the Royal Burgers’ Zoo, Netherlands. P. paniscus credit to William H. Calvin (https://commons.wikimedia.org/wiki/File:Bonobos_11yr_male_3yr_male_grin_Twycross.jpg)

The ‘cooperative eye hypothesis’ (6), suggests that the human depigmented sclera functions to facilitate hyper-cooperative behaviors (2, 6, 8) through, for example, the establishment of joint attentional states. That is, the ability of two or more individuals to jointly focus on one object or concept (6). A cooperation experiment, for example, found that participants who could view their partner’s gaze were better able to communicate referential meaning (about both the current and required location of objects) than where their gazes were obscured (9). A white background may signal iridal direction more conspicuously and in a manner that is more difficult to conceal than would a dark background (6, 10). Other primates can also follow conspecific gaze direction (11-14), but it is not yet clear whether they can follow gaze with a similar aptitude as humans. Only humans, for example, prioritize the use of eye direction over head direction when following a conspecific’s gaze (7). Humans can also determine conspecific iridal direction from a distance of up to 15 meters, though this may be partially mediated by the eyebrows (15). The cooperative eye hypothesis proposes that both the unusually depigmented human sclera and the potentially unusual sophistication of human gaze following could be associated with directional selection for cooperation.

Alternatively, the ‘self-domestication hypothesis’ posits that white sclerae may be pigment-related by-products associated with the human self-domestication process (7). Selection for tolerance and against aggression is predicted to result in the reduction in number and migration velocity of neural crest cells in early embryogenesis. This alteration may be responsible for domestication-syndrome - a range of behavioral and morphological traits that co-emerge with docility (16).

Pigment-producing melanocytes are derived from neural crest cells, so the reduction of melanocytes in a pale sclera (7) is potentially explicable as a correlated by-product of selection for social tolerance rather than as an explicitly functional adaptation.

The hypotheses describe above focus on the evolution of the human ocular morphology and the supposed uniqueness of human white sclerae. However, gaze cues are used widely in the primate order (e.g., gaze following or aversion; 11-14, 17). The ‘gaze camouflage hypothesis’, proposes that dark scleral pigment in primates may function as gaze direction camouflage from competitive conspecifics and predators (2). The ability to observe a conspecific’s gaze direction and, using that information and context, to learn about the emotional and/or intentional state of others may be a useful skill for organisms living within a social group (18-19). However, the act of observing another’s gaze direction may produce a source of information conflict. For instance, although it may be useful to third-party onlookers to receive social information (intentionally signaled or otherwise) as to one’s emotional and/or intentional states, it is not necessarily advantageous to the individual themselves to be sharing such information indiscriminately since this could be adversely utilized by competitors. For example, an experiment has been reported in which the location of food is revealed to a chimpanzee (witness), with another chimpanzee (witness-of-witness) unable to see the food’s location but able to observe the witness. The witness, potentially aware of having been observed, repeatedly misled the witness-of-witness by leading him to empty containers (20). Similarly, chimpanzees have been observed to avert their gaze from a high value food item if they, alone, are knowledgeable about its location and are in the presence of a dominant conspecific (17). These examples illustrate that communicating one’s visual direction conspicuously and/or indiscriminately may not always be an optimal social strategy.

Here, we utilize the remarkable diversity of primate ocular morphologies to provide the first quantitative, comparative investigation of the role of sociality in primate scleral color evolution. We compare scleral brightness with three behavioral measures: ‘proactive prosociality’, ‘social tolerance’, and ‘conspecific lethal violence’. Prosociality refers to any behavior which benefits another organism but where the originator does not benefit themselves (21). Prosociality can be reactive, e.g., in response to help-calls or subtle coercion (22), or proactive, that is, unsolicited (21). In humans, our heightened prosociality is considered to have facilitated the emergence of interdependent groups (23) as well as cumulative culture and technology (8). A previous report found the degree of allomaternal care to best predict observed patterns of proactive prosociality in primates, suggesting that increased prosocial tendencies may emerge with cooperative breeding systems (8). Social tolerance, that is, the tolerance towards groupmates and their interests, is not prosocial, but equally is also non-competitive. Meanwhile, competitive behaviors such as lethal violence are often similarly adaptive social strategies (24) as they may relate to territory or resource control (25) and can be flexible and coalitionary in format (26). These three measures enable the examination of the evolutionary role of scleral pigment across different social environment. Proactive prosociality is indicative of highly cooperative environments, social tolerance represents a behavior of intermediate social value and, by contrast, conspecific lethal violence is indicative of highly competitive social environments.

Under the gaze camouflage hypothesis, we therefore predict heightened conspecific lethal aggression to predict darker sclerae, and for dark sclerae to be likewise associated with low values of proactive prosociality and social tolerance. Both the cooperative eye hypothesis (directional selection) and the self-domestication hypothesis (correlated by-product) predict white sclerae to be associated with heightened proactive prosociality, although the self-domestication hypothesis explains the presence of prosociality as a correlated by-product of selection for social tolerance (8, 27-28). Hence, an association between scleral brightness and proactive prosociality, but not social tolerance, could be taken as evidence against the self-domestication hypothesis.

## Results

### Proactive Prosociality

Log (Scleral brightness) was significantly positively associated with sqrt (proactive prosociality) scores across the 15 primate species. Where lambda was taken at its maximum likelihood (λ_ML_=0), a statistically significant relationship was observed (p= < 0.001, R^2^=0.711, estimate=0.085, t=5.66; **Figure 2**). Similarly, where lambda was assumed to equal 1, a highly conservative comparison, the statistically significant relationship was still observed (p= < 0.001, R^2^=0.723, estimate=0.083, t=5.83).

**Figure 2.**
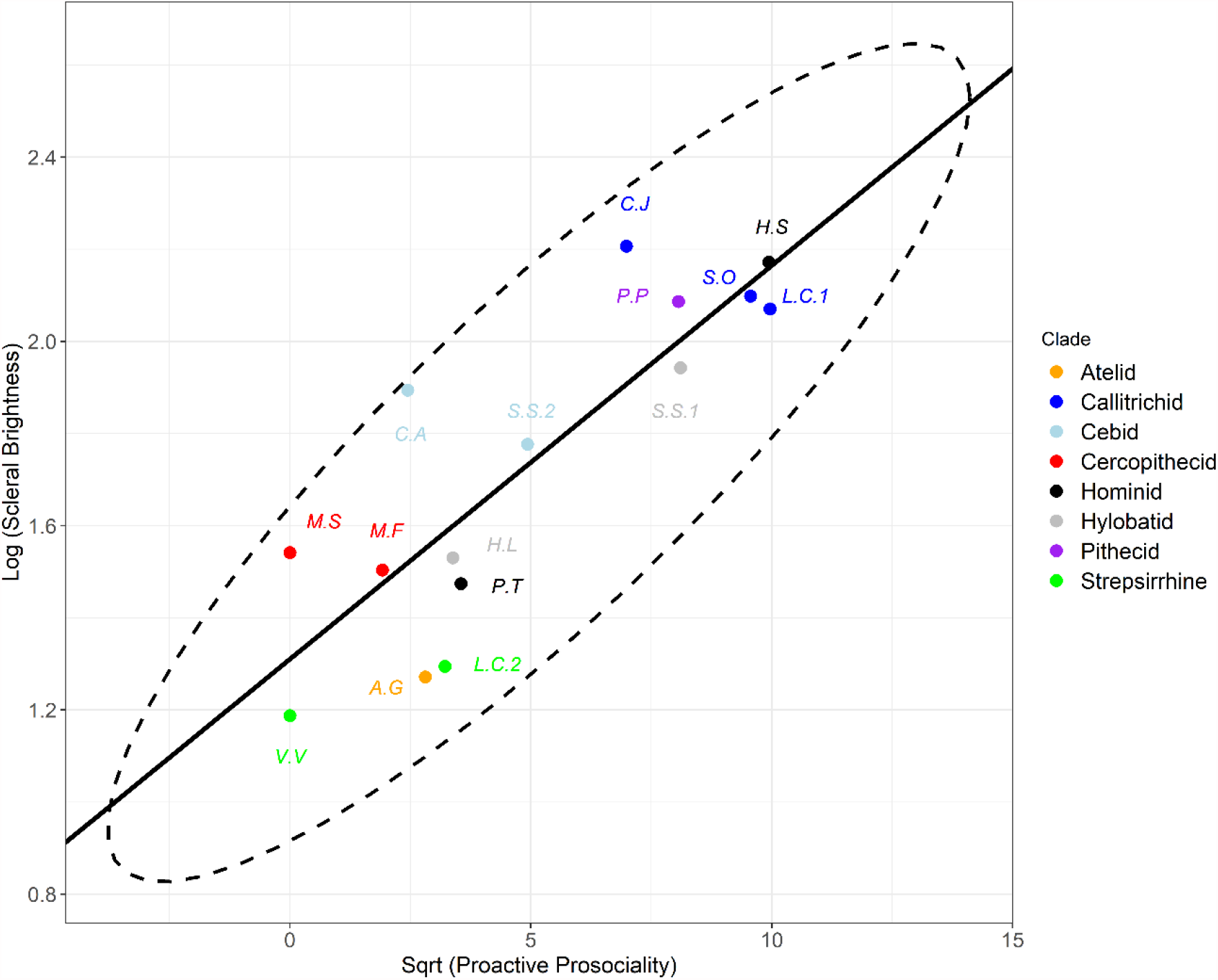
PGLS regression plot comparing log (scleral brightness) with sqrt (proactive prosociality) with in n=15 primate species. Confidence ellipse computed to 95% confidence. CJ = Callithrix jacchus; HS = Homo sapiens; SO = Saguinus oedipus; LC1 = Leontopithecus chrysomelas; PP = Pithecia pithecia; SS1 = Symphalangus syndactylus; SS2 = Saimiri sciureus; CA = Cebus apella; MS = Macaca silenus; MF = Macaca fuscata; HL = Hylobates lar; PT = Pan troglodytes; LC2 = Lemur catta; AG = Ateles geoffroyi; VV = Varecia variegata.

### Social Tolerance

Scleral brightness was significantly positively associated with social tolerance scores across the 15 primate species. Where lambda equals its maximum likelihood (λ_ML_=0), a significant relationship was observed (p=0.047, R^2^=0.269, estimate=90.58, t=2.19; **Figure 3**). Likewise, where weighted by lambda=1, the positive association between scleral brightness and social tolerance is still statistically significant (p=0.03, R^2^=0.314, estimate=95.57, t=2.44).

**Figure 3.**
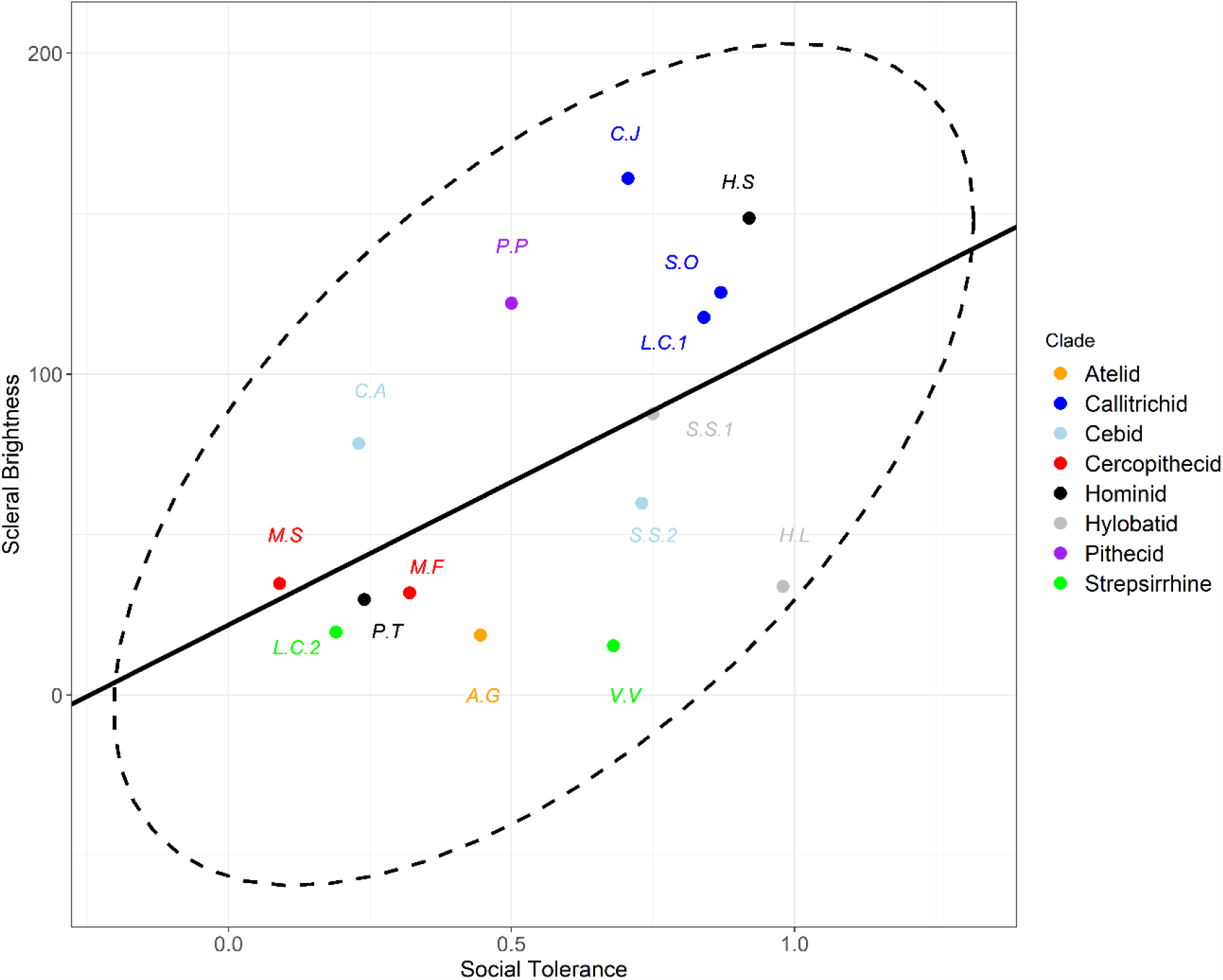
PGLS regression plot comparing scleral brightness with social tolerance scores in n=15 primate species. Confidence ellipse computed to 95% confidence. CJ = Callithrix jacchus; HS = Homo sapiens; SO = Saguinus oedipus; LC1 = Leontopithecus chrysomelas; PP = Pithecia pithecia; SS1 = Symphalangus syndactylus; SS2 = Saimiri sciureus; CA = Cebus apella; MS = Macaca silenus; MF = Macaca fuscata; HL = Hylobates lar; PT = Pan troglodytes; LC2 = Lemur catta; AG = Ateles geoffroyi; VV = Varecia variegata.

### Conspecific Lethal Violence

A significant negative association was observed across the 108 primate species between log (scleral brightness) and sqrt (conspecific lethal violence). Where lambda was taken at its maximum likelihood (λ_ML_=0.698), a statistically significant observation was observed (p= < 0.001, R^2^=0.118, estimate=-0.088, t=-3.77; **Figure 4**). Similarly, where lambda was assumed to equal 1, a statistically significantly relationship was still observed (p= < 0.001, R^2^=0.306, estimate=-0.144, t=-6.84).

**Figure 4.**
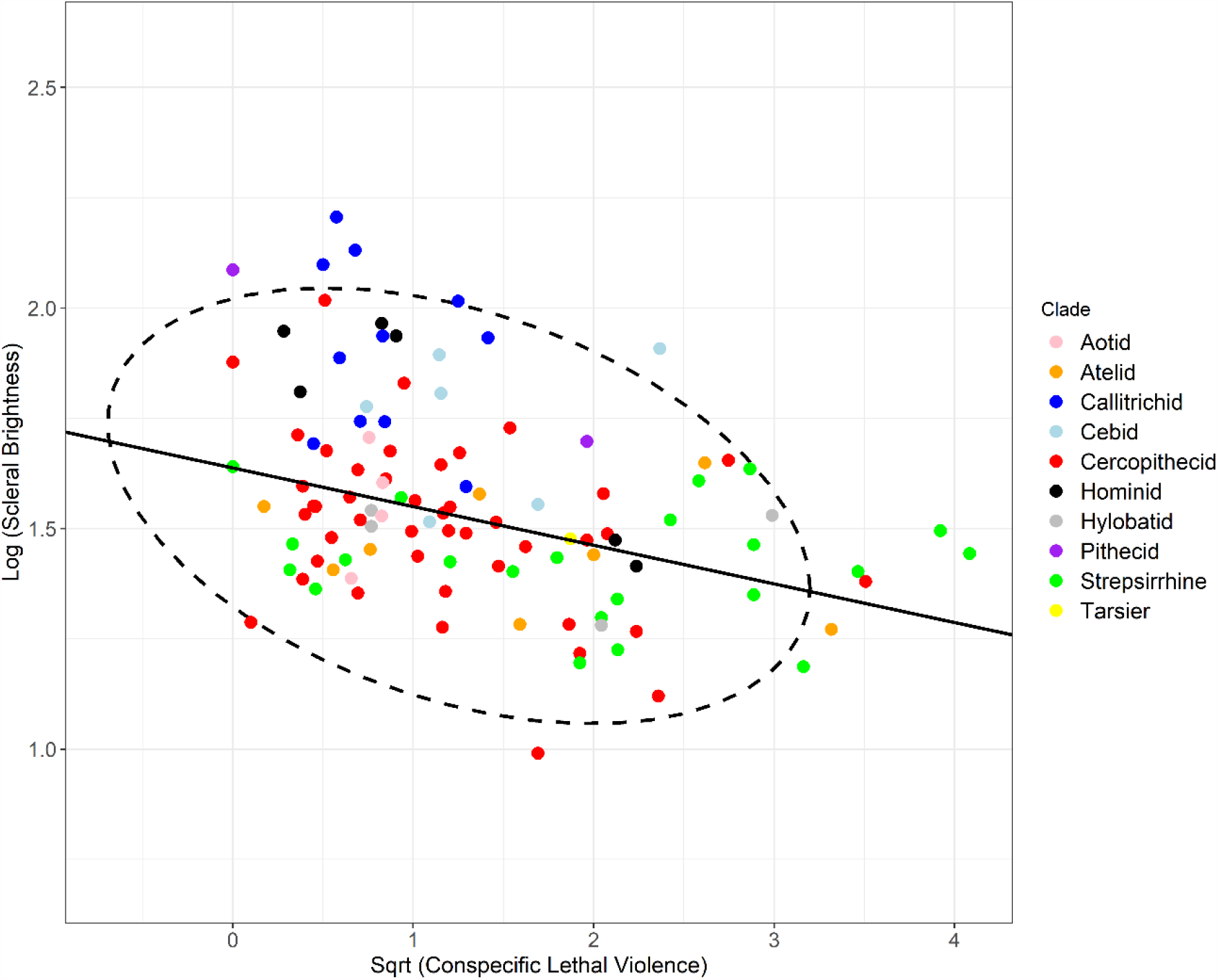
PGLS regression plot comparing log (scleral brightness) with sqrt (conspecific lethal violence). That is, the percentage of deaths due to conspecific lethal violence in n=108 primate species. Confidence ellipse computed to 95% confidence indicating bivariate outliers.

## Discussion

Our findings provide broad support for all three hypotheses tested: the gaze camouflage, cooperative eye and self-domestication hypotheses. We show that scleral pigmentation varies with differences in social behaviors between species of extant primates. Proactive prosociality, an experimentally derived measure of the degree to which individuals were willing to help their groupmates with no possibility of directly benefitting themselves, was associated with significantly increased scleral brightness (white sclerae being depigmented). Social tolerance, a measure of the evenness of the distribution of food items in the same species, was also associated with increased scleral brightness, though based on p-values and coefficients of determination (R^2^) appears to be a weaker predictor than proactive prosociality. Scleral brightness was significantly negatively associated with the percentage of deaths attributable to conspecific lethal aggression.

The presence of scleral pigmentation is likely to be a functional adaptation, rather than a product of random drift, due to the metabolic cost incurred in synthesizing dark pigment (2, 7). That the extent of conspecific lethal violence negatively predicts scleral brightness (i.e., predicts darker, more pigmented sclerae), may indicate that scleral pigment functions as a mechanism of gaze camouflage to conspecifics and/or predators (2). Chimpanzees, who are considered more reactively aggressive and less cooperative than humans and bonobos (7, 27), and who likewise possess significantly darker sclerae (5), have been shown to use visual concealment (29) and gaze aversion (17) when engaged in food competition. The presence of pigment could confer an advantage in terms of concealing gaze direction from groupmates and/or predators (though the latter was not tested here) in a competitive context (2). The fact that we have observed this relationship across the primate order is consistent with the conclusion that gaze following behaviors are widespread beyond the immediate hominid family (12).

Similarly, the presence of increased scleral brightness could also be adaptive – a position consistent with the cooperative eye hypothesis. Our results indicate that increased scleral brightness is associated with increased cooperative behaviors and reduced lethal violence. This finding is consistent with several previous studies. For example, a previous report found the degree of allomaternal care to best predict variation in proactive prosociality, arguing that this behavior may emerge with cooperative breeding systems (8). Additionally, alloparental care frequencies have been linked to neural control of facial musculature in primates (30), suggesting that the proper conveyance of non-verbal signals is of increased importance in cooperative breeding species. Furthermore, common marmosets (*Callithrix jacchus*), but not capuchin (*Cebus apella*) or tonkean monkeys (*Macaca tonkeana*), have been shown to share direct gaze when working on cooperative tasks that may be ambiguous (31-32). Of interest, these studies are also consistent with our finding that the four brightest species’ sclerae across our sample are found among humans and cooperative breeders: common marmosets (*Callithrix jacchus*), Goeldi’s marmosets (*Callimico goeldii*), humans and cotton-top tamarins (*Saguinus oedipus*). This finding is contrary to a commonly held belief that white sclerae are a uniquely human morphology (1-2, 6-7).

Alternatively, it may be that white sclera simply represent the absence of pigment which is synthesized only for use in competitive environments or, alternatively, could be by-products associated with self-domestication processes (7, 28). The self-domestication hypothesis predicts that selection for social tolerance, and against aggression, may generate prosocial behavior as a correlated by-product (27-28). Therefore, our finding that social tolerance, in addition to proactive prosociality, significantly predicts scleral brightness is consistent with the self-domestication perspective. However, we therefore cannot differentiate between self-domestication and cooperative eye explanations at this time. Furthermore, these may not be mutually exclusive perspectives. It could be, as an example, that white sclerae originated as a correlated by-product of self-domestication processes but subsequently became the subject of directional selection, thereby increasing fitness, as neural bases for social cue recognition have developed around this morphology (see 33).

These results provide a potential function and/or origin of divergent scleral morphologies and affirm that differences in scleral morphology are associated with differences in species’ social behavior. Recently, an investigation using relative measures (i.e., producing a ratio of the brightness contrast between the iris and sclera) found humans, bonobos and chimpanzees to have comparable, not distinct, gaze conspicuousness, despite their different scleral morphologies (5). This methodology, however, produces contrast ratios which may not be appropriate for large scale comparisons and which do not account for the physical properties of visible light that may bias the naturalistic perception of shade (34). Our results refute these findings and show that differences in primate scleral colorations are explicable with divergent social behaviors, indicating that the naturalistic perception of shade cannot be adequately captured with relative methodologies alone (34).

The sample sizes for the behavioral measures proactive prosociality and social tolerance may be considered small (both n=15). However, both represent the high amount of labor, hours and logistics required to experimentally extract these data (8). These behavioral data enable a deeper analysis than the use of common sociality proxies such as social group size. For example, many species, such as humans, chimpanzees and bonobos, live in large social groups (3) but differ widely in the social behaviors that are associated with ocular morphologies (7; 27). The use of such species-level behavioral proxies are therefore more appropriate, though the smaller sample size (and the use of different species within datasets) does exclude the possibility of multiple regression techniques. Likewise, the use of conspecific lethal violence data (35) acts as a useful indicator of a species’ aggressiveness. That said, the original compilation method did not separate between inter- and intra-group aggression, infanticide or maternal abandonment, behaviors that likely have different neural and functional bases (27) and which may exert different influences on scleral morphology. This may explain the weaker model fit in the conspecific lethal violence regression.

A limitation relates to the lighting properties of online photographs. Quantitative analyses of color from digital photographs often require photographs to be taken in controlled settings and/or with the use of a color standard for calibration (36). This is not possible when analyzing existing, uncalibrated photographs, however, this is less relevant when exclusively analyzing brightness rather than the additional hue and saturation values that comprise the full perception of color (rather than shade alone) from digital sources (34, 37). Although the ambient brightness of the photo represents a minor source of unaccounted variation, this is mitigated by the use of multiple distinct photos per species (minimum 6) with the majority of photos (97.6%) containing both eyes per individual for further average calculations. In this respect, we follow a previously established data collection methodology (5, 37). Furthermore, this approach, rather than a laboratory-based analysis, relates more closely to the naturalistic perception of ocular morphologies by onlookers since primates will continue to interact with conspecifics across a range of locations and times with different ambient shades. Hence, this enables more ecologically valid testing of the function of sclerae in relation to social interactions and allows us to examine a much larger dataset than would otherwise be possible.

In sum, we provide the first quantitative comparative analysis of the relationship between primate ocular pigment and social behavior. Human white eyes have long been a mystery by comparison to the dark scleral phenotypes observed among primates. Here, we show that pale scleral color is associated with cooperation whereas dark scleral color is associated with reduced cooperation and increased lethal aggression in extant primates. This refutes the recent notion that gaze conspicuousness be considered comparable, not distinct, between humans and *Pan* and lends support to the cooperative eye, self-domestication and gaze camouflage hypotheses of eye-behavior co-evolution.

## Materials and Methods

We obtained species-level scores of proactive prosociality and social tolerance across 15 extant primate species from (8). Full methodological details can be found with the original paper. In brief, the authors used a group service apparatus (38) to measure proactive prosociality and social tolerance. Captive primates were habituated to the apparatus and taught the function of an accessible lever which moved a board containing food into reach. Food was placed on the board in two positions - one where the individual pulling the lever could reach the food themselves and another where the individual pulling the lever could not reach the food themselves, but could make the food accessible to their group-mates. The resultant data are comparable between groups and species due to the standardization of this procedure. Proactive prosociality measured how many items of food an individual made available to their groupmates that they themselves could not access. Social tolerance was quantified where the board was in a fixed position with the food accessible and repeatedly replenished (35 times) as it was eaten. The authors then measured the evenness of the distribution of food items within the group to produce social tolerance scores. Lastly, conspecific lethal aggression was scored as the percentage of deaths per species (study populations are taken to be representative of each species) that were attributable to conspecific lethal aggression obtained from published data for 108 extant primate species (35).

We collected primate facial images from an online google image search completed in February 2021 and which used the common species name and the word “face” as key words. Some chimpanzee and bonobo images were collected from a previous study (5) and the Royal Burgers’ Zoo, Netherlands. Human images are public domain and were collected from the Pexels.com database using the key words “man”, “woman”, “face” and “eyes”. We then analyzed scleral brightness using ImageJ 1.x (39). Images were selected from this search to ensure the resolution of the external eye was of sufficient quality, that the eyes were unobscured by other objects and that there was no apparent photo manipulation present. Images were converted to greyscale such that the value of each pixel varied from 0 (black) to 255 (white) with intermediate scores being corresponding shades of grey. We then extracted scleral brightness values following existing techniques (5, 37). We collected values from within a rectangular selection area placed on the visible sclera for each eye per photo. A minimum of six photos were collected per species, although it is not known whether these represent six distinct individuals in all cases. We then calculated the median pixel value per selection area (due to the potential for outliers such as poor lighting, camera quality or light reflections) and the mean value per individual (i.e., mean of both eyes/selection areas), and subsequently the mean scleral brightness value per species. Of the 944 total facial photos, 21 (2.2%) presented with a cranial angle from which only 1 eye was clearly visible. In these few cases, the one accessible eye was taken to be representative of the scleral brightness of that individual. Species which were documented by (35), but for which insufficient number or quality of facial photos was available from online materials or which were not uniquely represented in the GenBank taxonomy were not included in analyses but are listed (see supplementary **Table S1**).

Phylogenetic generalized least squares analysis (PGLS) was completed in R version 4.0.3. (40) using the ‘caper’ package version 1.0.1. (41). A consensus phylogeny with branch lengths proportional to time (i.e., a chronogram) was generated and pruned for use from 10ktrees.com version 3 (42) and used the GenBank taxonomy (43). Variables were log transformed where residuals were non-normally distributed or to improve linearity. Where a variable contained zero-values, it was instead square root transformed as, unlike with log transformations, this does not require the input of an arbitrary constant which influences goodness-of-fit (44). Diagnostic plots including Q-Q, density and fitted and residual value plots were generated and inspected using the ‘plot.pgls’ function in the ‘caper’ package (41). Residual normality was further established using Shapiro-Wilk tests on model residuals. The significance value (i.e., alpha) is placed at 0.05. We estimated phylogenetic signal in PGLS analyses using Pagel’s λ, which indicates the degree to which the co-variance in model residuals is proportional to shared evolutionary history between species, assuming a Brownian motion model of evolutionary change over time (45-46). λ varies from 0 to 1, where 0 indicates that species are independent of one another and 1 the maximum level of phylogenetic signal, i.e., that co-variances are directly proportional to shared evolutionary history (45-46). Estimates of the maximum likelihood of lambda were subject to wide confidence intervals, a limitation increasingly common with reduced sample sizes (47) (likelihood profiles with confidence intervals are given in supplementary materials; **figures S1-S3**). For this reason, and to present a full picture of results, two result statements are provided per statistical test: one where lambda is assumed to equal its maximum likelihood (λ_ML_) and one where lambda is taken to equal 1 (the strictest phylogenetic control). The latter approach is highly conservative as higher phylogenetic control typically reduces the significance of independent variables (48).

## Acknowledgments

We would like Prof. Richard Wrangham for his comments on an earlier draft and the Royal Burgers’ Zoo, Netherlands for providing photo data.

