## Supplementary Materials for "The evolutionary origins of primate scleral coloration"

Electronic Supplementary Materials

| *Species* |
|---|
| *Callicebus hoffmannsi* |
| *Callicebus torquatus* |
| *Cercopithecus erythrotis* |
| *Cercopithecus pogonias* |
| *Cercopithecus preussi* |
| *Chlorocebus tantalus* |
| *Colobus satanas* |
| *Loris nycticeboides* |
| *Macaca sylvanus* |
| *Mandrillus leucophaeus* |
| *Mirza coquereli* |
| *Nomascus concolor* |
| *Nomascus siki* |
| *Oreonax flavicauda* |
| *Papio papio* |
| *Piliocolobus rufomitratus* |
| *Piliocolobus tephrosceles* |
| *Pithecia irrorata* |
| *Procolobus verus* |
| *Propithecus verreauxi* |
| *Saguinus nigricollis* |
| *Semnopithecus ajax* |
| *Tarsius bancanus* |
| *Tarsius spectrum* |
| *Trachypithecus delacouri* |
| *Trachypithecus poliocephalus* |
| *Brachyteles hypoxanthus* |
| *Cebus nigritus* |
| *Nomascus hainanus* |
| *Eulemur rufifrons* |
| *Pithecia monachus* |
| *Presbytis thomasi* |
| *Semnopithecus priam* |
| *Tarsius pumilus* |
| *Eulemur rufifrons* |
| Colobus vellerosus |
| Hylobates muelleri |
| Otolemur crassicaudatus |
| Presbytis chrysomelas |
| Procolobus verus |
| Saguinus leucopus |

| |
|---|
| Saguinus niger |
| Trachypithecus vetulus |
| *Cacajao melanocephalus* |

***Table S1****: List of species that were not included in analyses due to lack of sufficient quantity or quality of facial photo data or lack of representation in the GenBank taxonomy.*

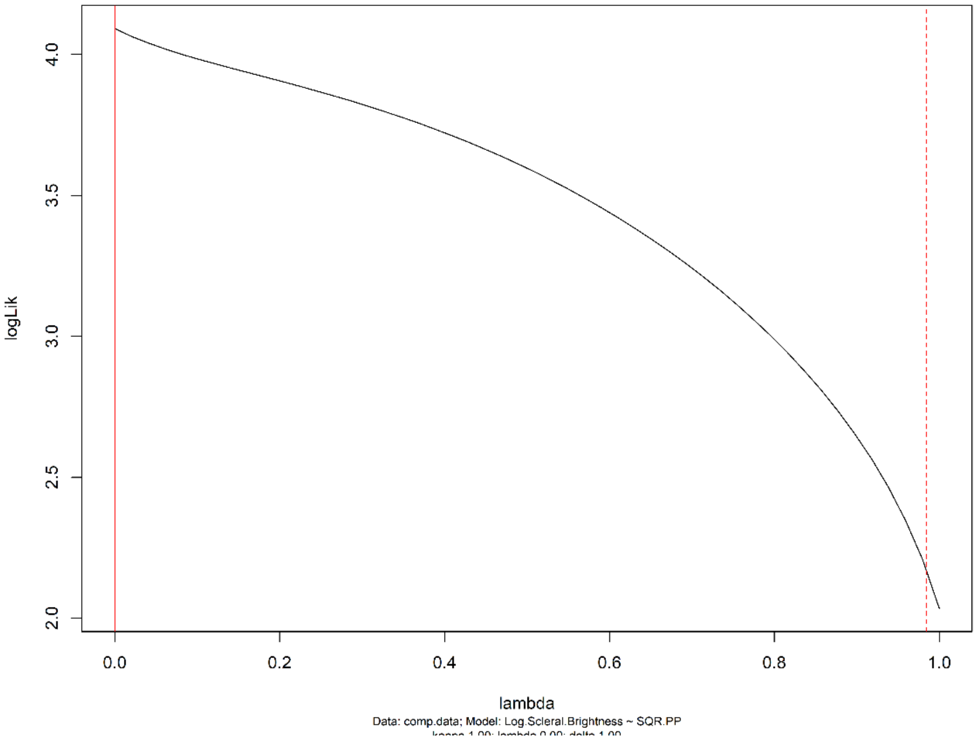


***Figure S1****: A profile showing the confidence intervals of lambda for the log (scleral brightness) ~ sqrt (proactive prosociality) PGLS regression. 95% confidence interval: 0-0.984. Due to wide confidence intervals, a conservative approach was used setting lambda at both maximum likelihood and 1.*

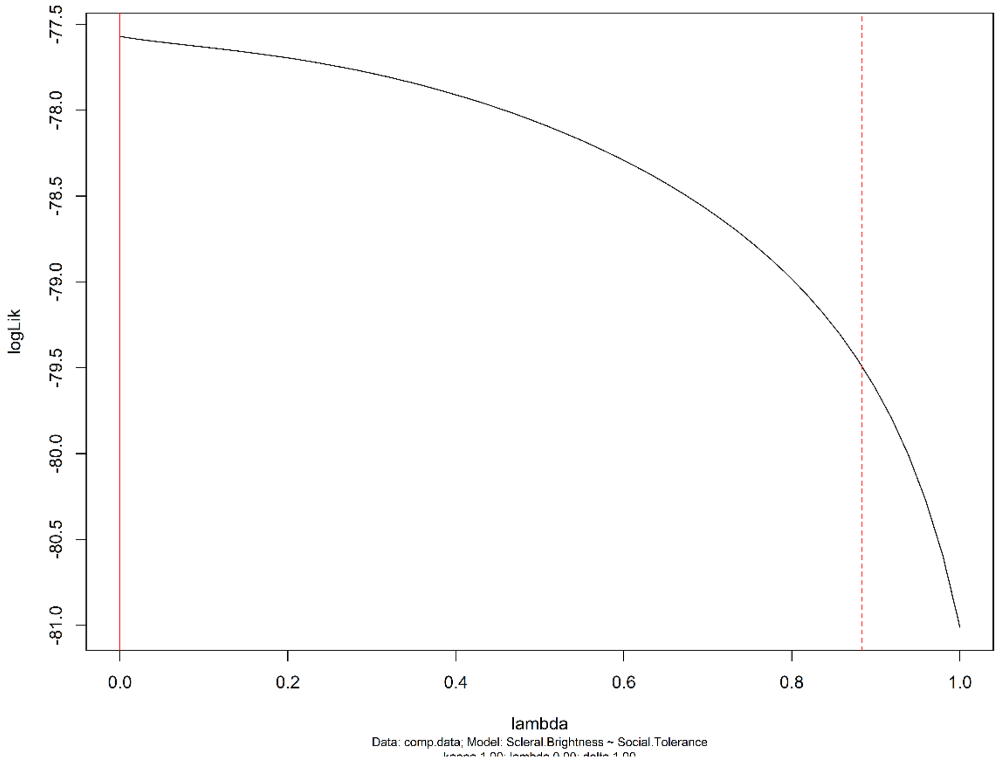


***Figure S2**: A profile showing the confidence intervals of lambda for the scleral brightness ~ social tolerance PGLS regression. 95% confidence interval: 0-0.883. Due to wide confidence intervals, a conservative approach was used setting lambda at both maximum likelihood and 1.*

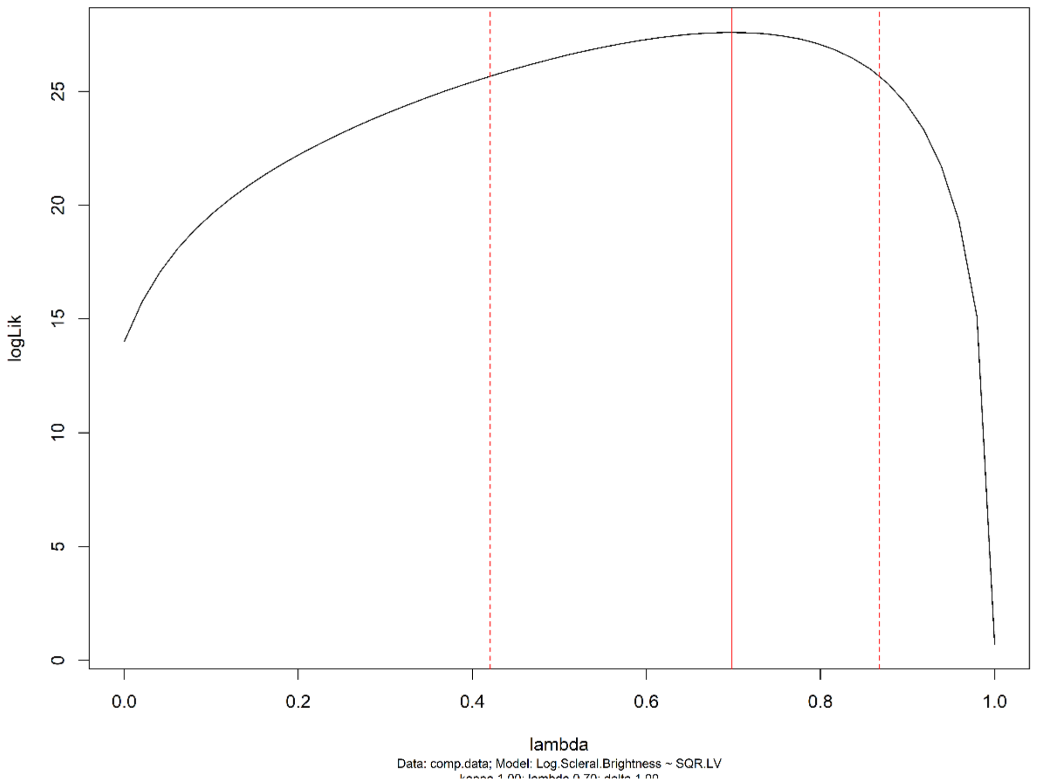

***Figure S3****: A profile showing the confidence intervals of lambda for the log (scleral brightness) ~ sqrt (conspecific lethal violence) PGLS regression. 95% confidence interval:* 0.421-0.868*. Due to wide confidence intervals, a conservative approach was used setting lambda at both maximum likelihood and 1.*
